# cloudrnaSPAdes: Isoform assembly using bulk barcoded RNA sequencing data

**DOI:** 10.1101/2023.07.25.550587

**Authors:** Dmitry Meleshko, Andrey D. Prjbelski, Mikhail Raiko, Alexandru I. Tomescu, Hagen Tilgner, Iman Hajirasouliha

## Abstract

**Motivation:** Recent advancements in long-read RNA sequencing have enabled the examination of full-length isoforms, previously uncaptured by short-read sequencing methods. An alternative powerful method for studying isoforms is through the use of barcoded short-read RNA reads, for which a barcode indicates whether two short-reads arise from the same molecule or not. Such techniques included the 10x Genomics linked-read based SParse Isoform Sequencing (SPIso-seq), as well as Loop-Seq, or Tell-Seq. Some applications, such as novel-isoform discovery, require very high coverage. Obtaining high coverage using long reads can be difficult, making barcoded RNA-seq data a valuable alternative for this task. However, most annotation pipelines are not able to work with a set of short reads instead of a single transcript, also not able to work with coverage gaps within a molecule if any. In order to overcome this challenge, we present an RNA-seq assembler allowing the determination of the expressed isoform per barcode.

**Results:** In this paper, we present cloudrnaSPAdes, a tool for assembling full-length isoforms from barcoded RNA-seq linked-read data in a reference-free fashion. Evaluating it on simulated and real human data, we found that cloudrnaSPAdes accurately assembles isoforms, even for genes with high isoform diversity.

**Availability:** cloudrnaSPAdes is a feature release of a SPAdes assembler and available at https://cab.spbu.ru/software/cloudrnaspades/.

## 1 Introduction

The emergence of long-read sequencing technologies, such as PacBio and Oxford Nanopore (ONT), have made it possible to perform transcriptome analysis at the isoform level and accurately predict full-length novel transcripts (Pertea et al. (2015); Tang et al. (2020); Nip et al. (2020); Kuo et al. (2020); Prjibelski et al. (2023); Sharon et al. (2013); Tilgner et al. (2014); Au et al. (2013)). However, even with the latest advances these technologies may feature an elevated number of sequencing errors or low coverage insufficient to reliably discover low-expressed isoforms.

Barcoded short-read RNA sequencing, such as SParse Isoform Sequencing (SPIso-seq, Tilgner et al. (2018)), Loop-Seq (Callahan et al. (2021)), and Tell-Seq (Chen et al. (2020)), presents a viable alternative to long-read sequencing, in particular for applications that require high coverage, such as estimating transcript abundance or capturing rare isoforms. While performing conventional bulk RNA sequencing, these technologies also allow to detect barcode sequences, which are identical for all read pairs sequenced from the same mRNA molecule. Considering the vast space of possible barcodes, the probability of identical barcodes being attached to mRNAs transcribed the same gene is low. Therefore, barcoded RNA sequencing combines high sequencing accuracy with linkage information between distant parts of the molecule, thus providing the opportunity to accurately reconstruct full-length isoforms.

While pipelines for the reference-based isoform reconstruction from barcoded data were developed in the original studies (Loop-Seq, SPIso-seq), to the best of our knowledge, there is currently no *de novo* RNA assembler specifically designed for barcoded RNA reads. Although Loop-Seq pipeline involves assembly of individual *read clouds* (reads sharing identical barcode sequence) using SPAdes assembler (Prjibelski et al. (2020)), it can only be applied when the coverage within a cloud is sufficient, which is often not the case, particularly for the SPIso-seq data (Tilgner et al. (2018)). Thus, the straightforward assembly of each cloud separately can lead to fragmented assemblies. At the same time, genome assemblers for barcoded sequencing data (Bankevich and Pevzner (2016); Tolstoganov et al. (2019)) focus on restoring long DNA fragments rather than isoforms sequences with high similarity and thus are not directly applicable to the mentioned types of data.

In this paper, we present *cloudrnaSPAdes*, a novel tool for *de novo* assembly of full-length isoforms from barcoded RNA-seq data. It first constructs a single assembly graph using the entire set of input reads and further derives paths for each read cloud, closing gaps and fixing sequencing errors in the process. Results on simulated and real human data show that *cloudrnaSPAdes* is able to accurately reconstruct full-length transcript sequences from read clouds having coverage as low as 1x, including genes with dozens of different expressing isoforms. As *cloudrnaSPAdes* does not require a reference genome or a gene annotation, it may become a useful tool for research projects studying previously unsequenced species.

## 2 Implementation

The outline of cloudrnaSPAdes is presented in Figure 1. First, an assembly graph is constructed from all input reads combined. Once constructed, the graph undergoes a basic simplification procedure (Bankevich et al. (2012); Nurk et al. (2013)), which removes only edges with extremely low coverage to prune obvious sequencing errors. Further, cloudrnaSPAdes algorithm attempts to reconstruct transcript sequences for each barcode. For the simplicity of explanation, we assume that reads are already grouped by barcode sequences, and the processing is done barcode by barcode.

**Fig. 1:**
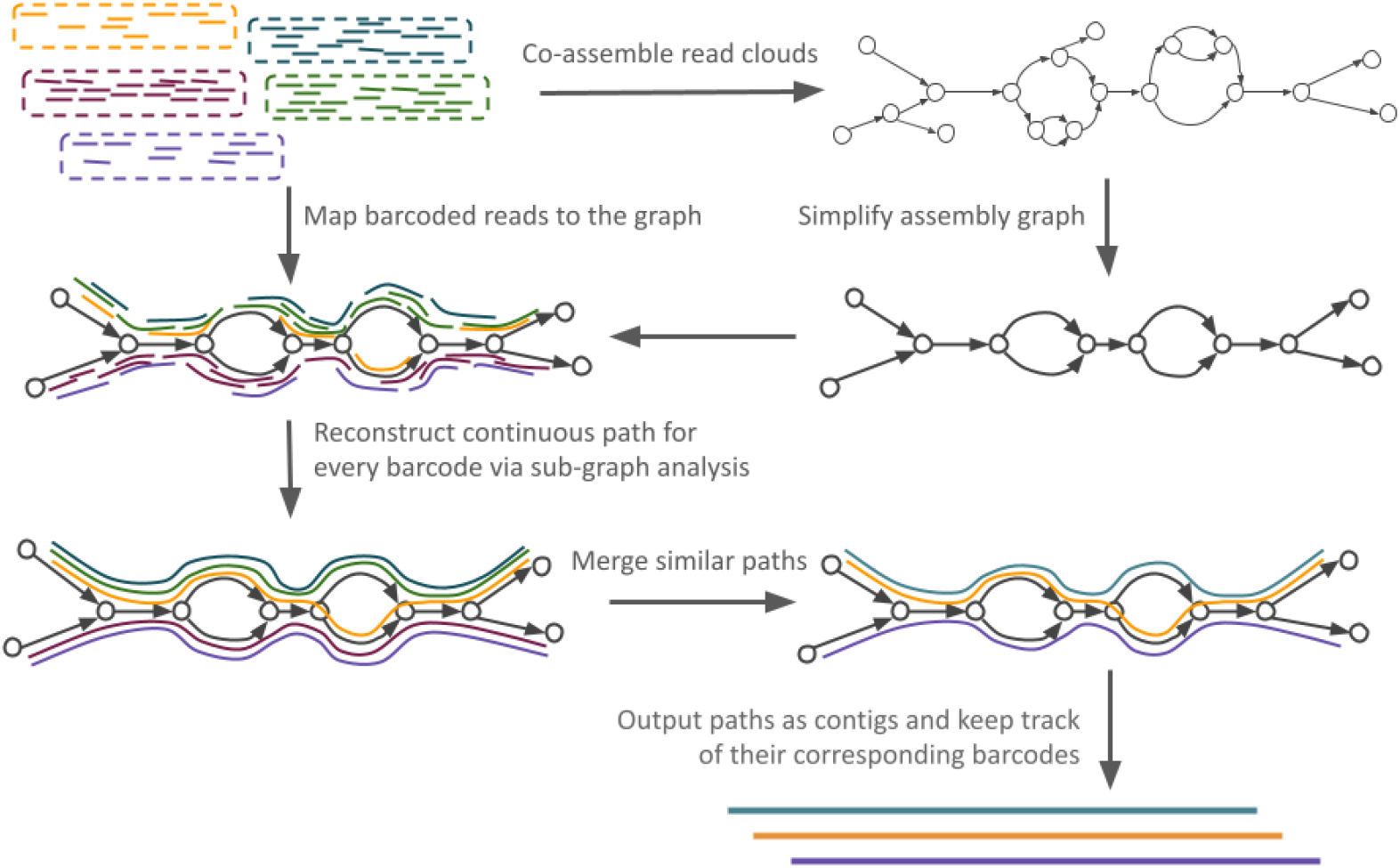
Outline of cloudrnaSPAdes. Initially, all reads are co-assembled into a single assembly graph. The graph is then simplified by removing low-covered tips and bulges. Subsequently, for each barcode, reads from the corresponding cloud are aligned to the graph and contigs are generated using obtained alignment information. Similar contigs are clustered together to calculate abundances and are outputted along with their corresponding barcodes.

The key intuition behind the algorithm is that different isoforms of the same gene are highly unlikely to be assigned an identical barcode. Thus, mapping reads from a read cloud to the assembly graph allows to detect non-overlapping paths corresponding to full-length transcript sequences. Once the reads from a read cloud are aligned to the assembly graph, we determine a subgraph formed by the alignments (i.e. a subset of edges that have at least one read mapped). Additionally, we compute barcode-specific coverage of the edges in the subgraph and determine the extreme positions (leftmost and rightmost) of the alignments for each edge in the subgraph.

The underlying assumption is that a set of reads with the same barcode originate from several mRNAs, and in an ideal scenario, the set of edges with alignments would form a subgraph consisting of multiple simple paths, where each path represents a single transcript. However, due to sequencing errors and coverage gaps, this assumption can be compromised. To address these challenges, we employ the following procedures in multiple cycles:

- **Tip clipping** - This step involves the removal of short tips (dead-end and dead start edges) and tips with low barcode-specific coverage from the subgraph.
- **Bulge Removal** - In this procedure, barcode-specific low-covered alternative paths are eliminated from the subgraph.
- **Gap closing** - Short edges from the assembly graph are added to the subgraph if they can be used to merge two simple paths into one, thereby closing gaps between them.

In comparison to conventional simplification procedures, cloudrnaSPAdes algorithm processes each read cloud individually and exploits barcode-specific edge coverage, while using the assembly graph constructed from all read clouds combined.

Once the subgraph is derived and simplified, we traverse this subgraph to extract a set of paths that represent isoforms. To accomplish this, we exploit exSPAnder algorithm (Prjibelski et al. (2014)), which utilizes linkage information from paired and single reads in order to reconstruct paths in the assembly graph. In this work exSPAnder was extensively modified to enable path construction using only a defined sub-graph and reads from a single cloud. The extracted paths are further clustered based on their sequence content, namely by their edges, as well as the leftmost position of the first edge and the rightmost position of the last edge they traverse.

Furthermore, we maintain a record of the barcodes associated with each cluster, enabling us to estimate the abundances of the respective isoforms. By storing this information, we can quantify the relative representation of different isoforms within the dataset, providing valuable insights into their prevalence and distribution.

Previous versions of exSPAnder were able to operate only on complete assembly graph, were not able to run multiple times in one assembly process, and ignored edges without paired-end information. By utilizing this modified version of the exSPAnder algorithm, we can effectively reconstruct isoforms, even in scenarios where they contain short repetitive regions within their sequences or when different isoforms of the same gene exhibit variations. This solution enables us to accurately restore the underlying structure of the isoforms, enhancing our ability to analyze and interpret complex genomic information.

The extracted paths are aggregated and organized within a data structure that groups them together based on the specific sequence of edges they traverse, as well as the leftmost position of the first edge and the rightmost position of the last edge they traverse. This approach is instrumental in distinguishing between isoforms that differ solely in their initial or final regions.

## 3 Results

### 3.1 Simulated data

To test the developed algorithms, we first selected three human genes: *GYPC, MAPT*, and *BIN1*, which exhibit varying levels of alternative splicing complexity (5, 16, and 15 isoforms respectively). For each gene, we generated simulated reads using InSilicoSeq (Gourlé et al. (2019)), ensuring that cloud sizes fit the distribution observed in real SPIso-seq data. We assembled reads for each gene using cloudrnaSPAdes, and then used IsoQuant (Prjibelski et al. (2023)) to assign each assembled contig to a reference isoform. For each barcode, we compared the assigned isoform with the isoform we used for simulation. As a competitor strategy, we implemented a strategy that assembles read clouds separately. The same strategy is implemented by LoopSeq Callahan et al. (2021), that use SPAdes with unspecified parameters for each subassembly. LoopSeq pipeline suggests that each read cloud is sequenced around 30*×*, so we suppose that default parameters were used, but for working with low-coverage data we recommend ‘–sc’ parameter as the most effective.

Table 1 demonstrates that cloudrnaSPAdes is capable of assembling isoforms, more than 98% of which are accurately assigned using IsoQuant. Importantly, we observed no consistent misclassification of any particular isoform. However, we did observe instances where read clouds were ambiguously or incorrectly classified, and the frequency of such occurrences increased for the read clouds with the lower number of reads. Therefore for such read clouds, the isoform is likely to be split into several parts due to coverage gaps. However, even in real data, we can identify such cases by examining whether the same read cloud produced multiple isoforms for a specific gene. This situation indicates that either the read cloud was not assembled as a single isoform or that two distinct isoforms appeared within the same read cloud. Both types of events should are not beneficial for subsequent downstream analysis and can be filtered. The “Fixed recall” column of Table 1 shows that after this procedure, the recall goes up for the complex MAPT and BIN1 genes. Separate strategy also showed high precision for GYPC and BIN1 genes, but recall is much lower compared to cloudrnaSPAdes.

**Table 1.** Quality assessment results on GYPC, MAPT, and BIN1 genes.

| Gene | Number of Barcodes | Number of Barcodes with $\geq 5$ read pairs | Precision cloudnaSPAdes | Recall cloudnaSPAdes | Fixed recall cloudnaSPAdes | Precision Separate | Recall Separate | Fixed recall Separate |
| --- | --- | --- | --- | --- | --- | --- | --- | --- |
| GYPC | 539 | 320 | 1.0 | 0.997 | 0.997 | 1.0 | 0.839 | 0.891 |
| MAPT | 1702 | 989 | 0.984 | 0.712 | 0.808 | 0.914 | 0.515 | 0.810 |
| BIN1 | 1399 | 840 | 0.991 | 0.773 | 0.831 | 1.0 | 0.564 | 0.722 |

We also assessed the performance of popular short-read RNA assemblers in the case of complex simulated genes. Standard RNA assemblers would assemble the whole dataset ignoring barcode information, and restore transcripts using paired-end information and graph topology. We can’t directly compare these assemblers with cloudrnaSPAdes and Separate pipeline, since they work on a per-barcode basis. However, we can count the number of full-length isoforms in the transcripts produced by various assemblers. We will count the isoform as restored if there is a unique assignment of the contig to the isoform in Isoquant. For our experiments, we ran the most popular RNA assemblers - Trinity Grabherr et al. (2011) and rnaSPAdes Bushmanova et al. (2019). Results can be found in Table 2. Note that cloudrnsSPAdes and “Separate” pipeline were able to identify each isoform multiple times, but for the standard short-read RNA assemblers most of the isoforms were undetected. Nevertheless, Trinity performs better than rnaSPAdes on our datasets.

**Table 2.**
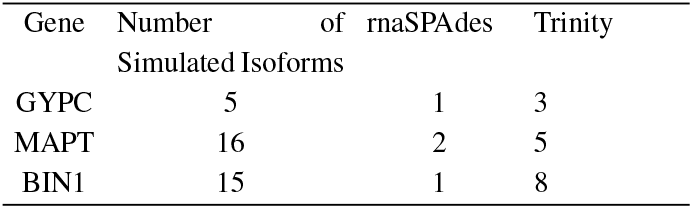
Quality assessment of short-read RNA assemblers results on GYPC, MAPT, and BIN1 genes. Both Trinity and rnaSPAdes are not able to restore the most of the isoforms.

### 3.2 Real data

To demonstrate the effectiveness of our method in detecting full-length isoforms, we conducted an analysis using SPIso-seq sequencing data obtained from a diverse pool of 50 individual human brain cDNA samples (Tilgner et al. (2018)). This dataset enabled us to explore a wide range of alternative isoforms and natural variations within genes. After aligning the reads to the human genome using STAR aligner (Dobin et al. (2013)), we focused specifically on the reads that mapped to chromosome X. From this subset, we successfully assembled a total of 6.8 million read pairs, corresponding to 245,098 unique barcodes, utilizing our cloudrnaSPAdes tool. The assembly process required approximately 9 hours and reached a peak memory usage of 14 GB.

The assembly process yielded a total of 89,108 contigs with a length greater than 300 bp. To gain insights into the isoform composition, we utilized IsoQuant in combination with GENCODE v32 annotation and the GRCh38 reference genome. Among these contigs, 22,936 exhibited a unique assignment to a specific isoform, while 4,040 contigs had minor differences in their assignments. In contrast, 34,553 contigs were assigned ambiguously, 12,404 contigs showed inconsistencies in their assignments, and 10,492 assignments were deemed uninformative.

The presence of a substantial number of non-unique assignments can be attributed to the inherent noise in real sequencing data. It is important to note that not only full-length isoforms are captured, but also fragmented or partially processed RNA products, which can contribute to the ambiguity in assignments. However, since processed isoforms tend to be more stable compared to other RNA states, we can leverage a barcode-based filtering strategy to mitigate this issue.

By utilizing read clouds, we can filter the contigs based on the number of associated barcodes. This approach is particularly effective in removing contigs that are classified as inconsistent, non-informative, or intergenic. Interestingly, the filtering method demonstrates similar removal rates for both ambiguously and uniquely assigned contigs. This suggests that at least some ambiguously classified contigs are produced by multiple read clouds. Various factors, such as incomplete annotation or unconnected regions in the assembly graph, could contribute to this phenomenon. A comprehensive summary of the filtering results is presented in Table 3.

**Table 3.** Isoquant assignment results on real dataset.

| Minimal number of barcodes | Unique | Unique with minor difference | Ambiguous | Inconsistent | Non-informative | Intergenic |
| --- | --- | --- | --- | --- | --- | --- |
| 1 | 22936 | 4040 | 34553 | 12404 | 10492 | 1699 |
| 2 | 8609 | 376 | 13773 | 1359 | 1287 | 267 |
| 3 | 4662 | 186 | 7845 | 438 | 240 | 41 |
| 4 | 3080 | 118 | 5290 | 273 | 65 | 13 |
| 1-fixed | 22719 | 1238 | 29837 | 11671 | 8473 | 1217 |
| 2-fixed | 6856 | 310 | 9923 | 1130 | 778 | 123 |
| 3-fixed | 3502 | 140 | 5171 | 357 | 169 | 21 |
| 4-fixed | 2228 | 82 | 3251 | 224 | 49 | 7 |

To further explore the effectiveness of filtering procedures in barcoded RNA assembly, we developed a filtering method similar to the procedure described in the “Simulated data” section. Specifically, we implemented a strategy to filter contigs, if they have a barcode that was assigned to a single gene multiple times. This approach aimed to remove parts of read clouds with low coverage for a specific gene, thereby enhancing the accuracy of the assembly results. The results are presented in Table 3 in “fixed rows”. A comparison between the “1” and “1-fixed” rows in the table reveals that approximately 15% of the ambiguous assignments can be effectively filtered out. This occurs when a single read cloud produces multiple contigs assigned to the same gene. On the other hand, the unique assignments remain relatively unchanged throughout the filtering procedure. These findings suggest that different strategies for refining and improving the results using barcode information hold promise for future investigations. This part highlights the potential of leveraging barcode-based filtering methods to enhance the accuracy and interpretability of barcoded RNA assembly. By refining the assembly results and reducing the presence of ambiguous assignments, researchers can gain more confidence in their analyses and further explore the complexity of isoform expression in various biological systems.

To demonstrate the effectiveness of cloudrnaSPAdes in reconstructing different isoforms of a single gene, we selected the ATP6AP2 gene from chromosome X. This gene plays a crucial role in normal kidney development and function Hoffmann and Peters (2021). As documented in GENCODE v32, it has a total of 34 distinct isoforms, although we do not expect all of them to be expressed in our dataset. Remarkably, cloudrnaSPAdes successfully assembled seven isoforms of ATP6AP2, highlighting its ability to capture and reconstruct different variants of a gene.

In contrast, a conventional *de novo* transcriptome assembler, rnaSPAdes Bushmanova et al. (2019), failed to restore any isoforms with unique assignments for this particular gene. This emphasizes the better performance of cloudrnaSPAdes and utility of read clouds in general in accurately reconstructing isoforms and obtaining reliable assignments. The alignments of some of the assembled isoforms are depicted in Figure 2, illustrating the diversity of isoforms captured by cloudrnaSPAdes.

**Fig. 2:**
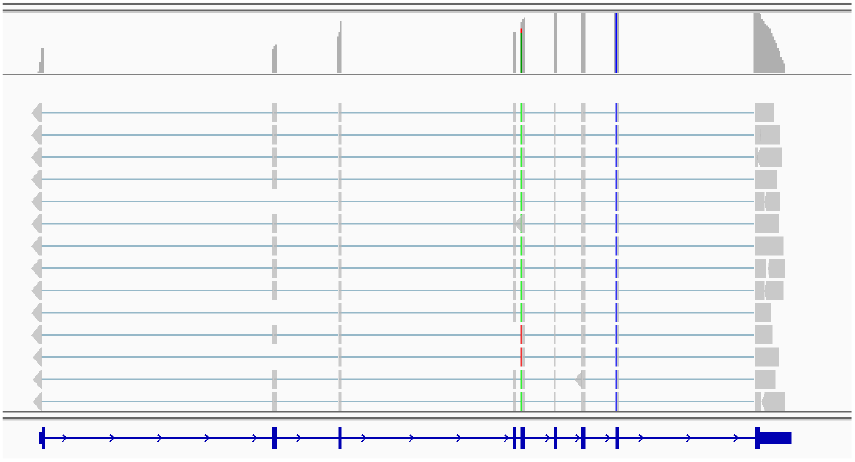
Few alignments of ATP6AP2 gene. Contigs were produced by cloudrnaSPAdes. Among these alignments we can see isoforms where second exon is missing, forth exon is missing, both alternative exons are missing, and both exons are present.

## 4 Discussion

The rise of long-read technologies, such as PacBio and Oxford Nanopore sequencing, has overshadowed the utility of read clouds in contemporary computational biology. These long-read technologies offer easier data interpretation and yield substantial results, capturing the attention of researchers across various computational biology domains.

Despite this trend, there remains a niche where read clouds can exhibit comparable performance to long reads and find success in the market. Specifically, in the realm of transcriptome sequencing, the relatively simpler nature of the data allows read clouds to achieve results of similar quality to long reads. Transcriptome analysis often demands extensive sequencing data to uncover rare isoforms, and the substantial input requirements and sequencing costs associated with long reads can make read clouds a compelling alternative.

Therefore read clouds can be effectively used for discovering novel isoforms and isoform discovery in non-human species. cloudrnaSPAdes offers an effective way to assemble full-length isoforms from cheap and accurate read cloud data. Uncovering novel isoforms plays a crucial role in advancing our understanding of gene regulation, alternative splicing patterns, and functional diversity across different organisms. By identifying and characterizing isoforms in non-human species, we gain insights into the evolutionary dynamics and adaptation processes specific to those organisms.

Notably, platforms like Tell-Seq and Loop-Seq continue to thrive in the market, specifically catering to transcriptomic analysis. These platforms recognize the unique advantages offered by read clouds in this particular context and continue to provide valuable solutions to researchers in need of comprehensive transcriptome analysis.

## Funding

This work was supported by the NIGMS Maximizing Investigators’ Research Award (MIRA) R35 GM138152 to I.H. M.R. was supported by St. Petersburg State University (grant ID PURE: 73023672).

### Conflicts of interest

none declared.

## Notes

### Competing Interest Statement

The authors have declared no competing interest.

